# DOT1L induces RNAPII accumulation independent of its catalytic activity in pluripotent stem cells

**DOI:** 10.1101/2025.08.08.668971

**Authors:** Xiaoya Zhang, Rupa Sridharan

**Affiliations:** Wisconsin Institute for Discovery, University of Wisconsin-Madison, Madison, WI 53715, USA; Laboratory of Genetics, University of Wisconsin-Madison, Madison, WI 53715, USA; Department of Cell and Regenerative Biology, University of Wisconsin-Madison, Madison, WI 53715, USA

## Abstract

Pluripotent stem cells can be propagated in vitro from embryos in specific culture conditions that capture subtle developmental transitions. Naïve pluripotent mouse embryonic stem cells (ESCs) can exist in two distinct epigenetic states—the metastable state in serum/LIF conditions and the ground state with pharmacological inhibition of differentiation-inducing pathways in the 2i/LIF conditions, which better resembles the in vivo blastocyst. Here, we acutely induced one feature of 2i/LIF ESCs, an increase in the H3K79 methyltransferase, DOT1L, in serum/LIF ESCs to determine its effects on metastable pluripotency. We find that DOT1L induction causes an increase in RNA Polymerase II (RNAPII) accumulation at the transcription start site, irrespective of catalytic activity, mimicking 2i/LIF RNAPII pattern. However, the pulse of DOT1L and consequent RNAPII accumulation is insufficient to induce immediate changes in steady-state or nascent RNA expression. Genes with higher transcription and elongation rates exhibit moderate changes in RNAPII accumulation, while lowly transcribed genes separate into two distinct groups, with one group showing the strongest RNAPII accumulation in response to DOT1L induction and the other showing the weakest. This differential accumulation is reduced at H3K27me3-enriched and bivalent genes. We also find that cells that sustain DOT1L expression have a homogenous NANOG protein profile without affecting *Nanog* transcription. Taken together, we find that a pulse of DOT1L in serum/LIF ESCs is sufficient to partially recapitulate certain features of ground state pluripotency, reinforcing its importance in this state of the pluripotency continuum.

## Introduction

During mouse embryonic development, pluripotency, the ability to give rise to cells from all three germ layers, is fleeting. It is restricted spatially to the inner cell mass of the blastocyst or the epiblast and temporally from ∼embryonic day (E) 3.5 to ∼ E5.5, respectively^1,2^. The pluripotency continuum refers to the transitions from a “naïve” state in the blastocyst to a “formative” state in the newly implanted embryo before acquiring a “primed” state as cells prepare for germ layer differentiation during gastrulation^3,4^. The inner cell mass can be isolated until about E4.5 and propagated in vitro to derive mouse embryonic stem cell (ESC) lines. These naïve ESCs, despite continuously self-renewing in vitro, can be injected in vivo into blastocysts and contribute to all ensuing germ layers and germ cells^5–7^.

Functionally naïve pluripotent ESCs can be derived in two distinct culture conditions that keep autocrine and paracrine differentiation cues from the FGF-ERK pathway at bay^8–10^. When ESC lines are derived in serum along with the cytokine leukemia inhibitory factor (LIF), the bone morphogenetic (BMP4) signaling in serum blocks ectoderm, while LIF blocks the endoderm and mesoderm differentiation^11–13^. Since these signals block differentiation to opposing cell fates, the serum/LIF established ESCs are considered metastable. Pluripotent ESCs can also be derived in the presence of two pharmacological inhibitors, 2i – to the MEK/ERK pathway (PD0325901), and the GSK3β (CHIR99021) that activates Wnt signaling^8,9^. Together with LIF, the 2i propagated ESCs are in a ground state that blocks the differentiation pathways^14^. Molecularly, the 2i/LIF derived ESCs cluster closely with E4.5 epiblast cells, both at the single-cell level, transcriptomic level, and in similarity of epigenetic features^13,15,16^.

Among transcriptional changes, in contrast to serum/LIF ESCs, 2i/LIF ESCs display reduced expression of lineage-associated genes, homogenous expression of pluripotency transcription factors such as *Nanog,* and lower expression of *c-Myc*^16,17^. 2i/LIF ESCs have globally lowered DNA methylation due to the increased expression of *Prdm14*^18,19^. At promoters of developmentally regulated genes, serum/LIF ESCs display bivalency with enrichment of opposing histone modifications- the activating H3K4me3 and repressive H3K27me3^16,20–22^. Bivalency is predominant at promoters of lineage-associated genes, which was thought to represent a poised chromatin state. This state is diminished in 2i/LIF ESCs because of a reduction in H3K4me3^22^. By contrast, Polycomb-mediated H3K27me3 in 2i/LIF ESCs is globally increased but redistributed away from bivalent promoters to intergenic regions, contributing to their DNA-hypomethylated epigenetic landscape^16,23^. Prior unbiased histone mass spectrometry data revealed that 2i/LIF ESCs had higher levels of H3K27me3 and H3K79me2, and lower levels of H3K27ac and H4 acetylation^23^. In addition to these epigenetic differences, there are also changes in transcriptional dynamics. Compared to serum/LIF ESCs, 2i/LIF ESCs exhibit increased RNA polymerase II (RNAPII) pausing, slower transcriptional elongation rates, and lower total transcript levels^16,24^. One contributing factor is the marked downregulation of MYC, which promotes transcriptional pause-release^16,25–27^.

Our prior work has indicated that DOT1L-mediated H3K79 methylation contributes to the hypertranscription phenotype of pluripotent stem cells and modulates RNAPII enrichment patterns during the acquisition of pluripotency from somatic cells^28^. Most existing studies have focused on loss-of-function approaches to examine DOT1L in ESCs^29–31^. However, it remains unclear how DOT1L abundance and H3K79me levels are regulated across the pluripotency continuum, and whether elevated H3K79me contributes to the unique epigenetic landscape and RNAPII dynamics of 2i/LIF ESCs. To address this, we performed acute overexpression of DOT1L in serum/LIF ESCs. We found that increased DOT1L causes an enrichment of RNAPII even without functional catalytic activity. The extent of RNAPII enrichment was generally proportional to absolute expression levels and RNAPII elongation rates in serum/LIF ESCs. By contrast, the group of genes that contained elevated H3K27me3 levels and that were overrepresented for bivalent genes did not gain much RNAPII. The acute increase in DOT1L also mimicked the homogenous expression of NANOG in 2i without an immediate effect on transcription. Elevated DOT1L levels naturally found in 2i conditions may be sufficient to alter RNAPII enrichment levels in the ground state of pluripotency at highly expressed genes.

## Results

### H3K79me2 increase in 2i/LIF ESC is a consequence of elevated DOT1L levels and slower RNAPII kinetics

We had previously found that H3K79 methylation levels were higher in 2i/LIF as compared to serum/LIF conditions^28^. This could indicate a higher protein abundance of DOT1L, the only H3K79 methyltransferase. RNAPII is known to recruit DOT1L, a distributive enzyme, where the higher local concentration would increase H3K79 methylation levels^32,33^. H3K79me could also accumulate in 2i/LIF ESC because of their slower RNAPII pause release and elongation rates. The slower RNAPII dynamics in 2i/LIF could result from lowered abundance of RNAPII, similar to our observation in somatic cells. To interrogate these possibilities, we first assessed the abundance of RNA Polymerase II (RNAPII) and the phosphorylation status of its C-terminal domain (CTD) in embryonic stem cells (ESCs) cultured in 2i/LIF (2i) versus serum/LIF (serum) conditions (Figure 1A). 2i/LIF ESCs displayed elevated levels of total RNAPII protein, along with increased Serine 5 phosphorylation (Ser5P), associated with transcription initiation, and Serine 2 phosphorylation (Ser2P), linked to elongation, when compared to serum/LIF ESCs (Figure 1A). Thus, unlike somatic cells, which have a lower level of RNAPII and slower transcriptional elongation rates as compared to serum/LIF ESCs, 2i/LIF ESCs, despite being slower, have a higher level of RNAPII^24,28^. Next, we tested the abundance of DOT1L protein and found that DOT1L levels are approximately 1.5-fold higher in 2i/LIF ESCs relative to serum conditions, contributing to the increase in H3K79me2 (Figure 1B).

**Figure 1.**
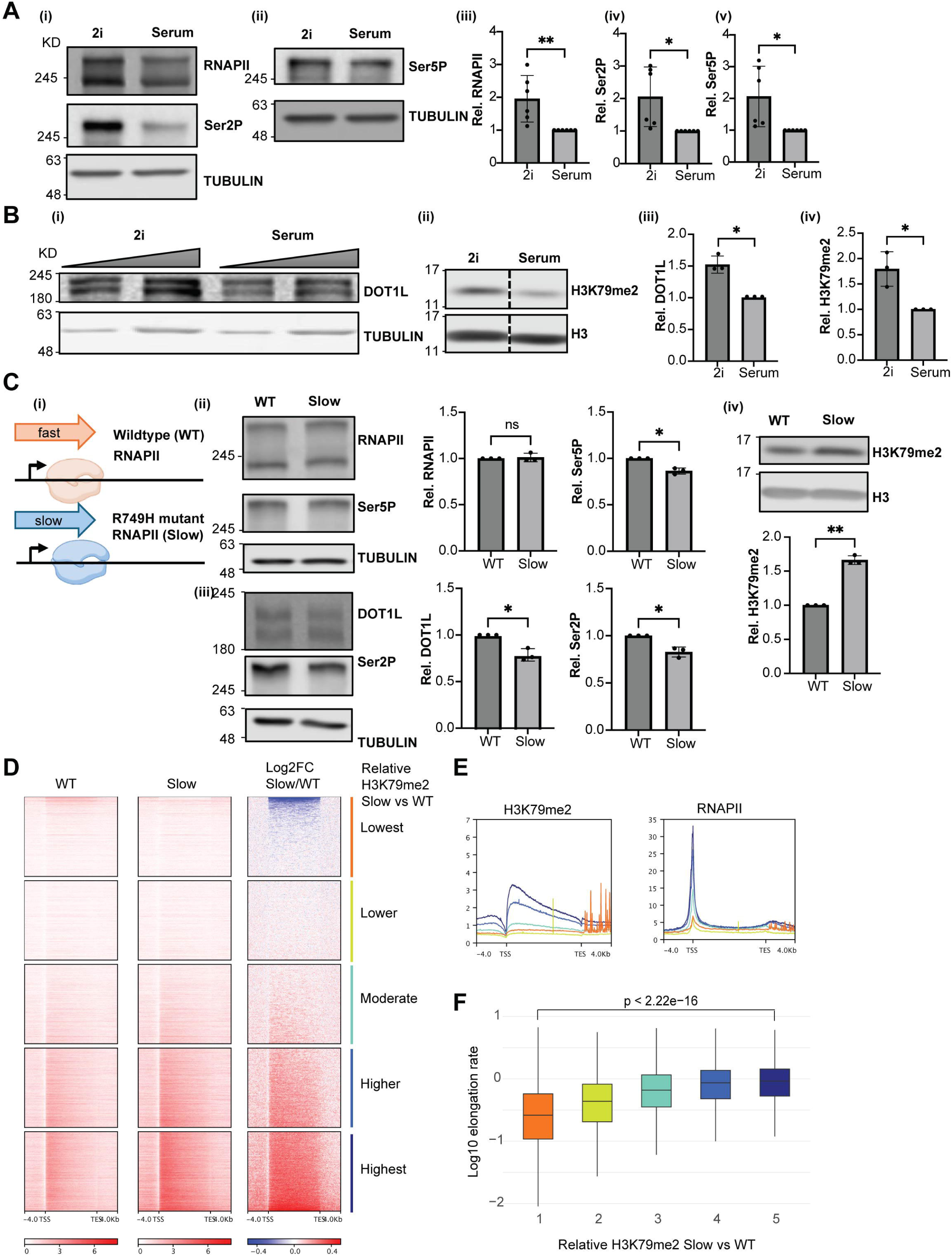
**RNAPII dynamics and H3K79me2 levels are altered between pluripotent states and modulated by transcription elongation rates.** (A) Comparison of RNAPII abundance and phosphorylation profiles in ESCs cultured in serum versus 2i.(i-ii) Western blot of total RNAPII protein levels and Ser2P (i) and Ser5P (ii) with TUBULIN loading control in 2i and serum LIF ESCs. (iii–v) Quantification of RNAPII Ser2P, Ser5P, and total RNAPII normalized to TUBULIN (mean ± SD, * = p < 0.05, ** = p < 0.01 by ratio paired two-tailed t test, n = 6). (B) (i) Western blot of DOT1L and TUBULIN control in serum versus 2i ESCs with increasing protein loading. (ii) Western blot of H3K79me2 and H3 loading control in 2i and serum LIF ESCs. Dashed line indicates cropped blot. (iii–iv) Quantification of DOT1L and H3K79me2 normalized to TUBULIN and total H3 respectively (mean ± SD, * = p < 0.05, ** = p < 0.01 by ratio paired two-tailed t test, n=3). (C) (i) Schematic of wildtype versus slow-mutant RNAPII. (ii–iii) Western blots of total RNAPII, Ser5P (ii), DOT1L, Ser2P (iii) with TUBULIN control in WT versus slow mutant ESCs. (iv) Western blot of H3K79me2 with H3 control in WT and slow ESCs. Graphs show quantification relative to tubulin or H3 (mean ± SD, * = p < 0.05, ns = not significant, by ratio paired two-tailed t test, n=3). (D) Heatmaps showing H3K79me2 ChIP-seq signal in WT and slow mutant (Slow) ESCs, sorted and grouped evenly by Log2FC slow vs WT into quintiles. (E) Metaplots of H3K79me2 in WT ESCs and Marks et al. RNAPII ChIP-seq in 2i/LIF ESCs enrichment of grouped by relative H3K79me2 slow/WT ratio in D (color matching)^16^ . (F) Boxplot of elongation rates in 2i/LIF ESCs for genes grouped by relative H3K79me2 in slow/WT ratio.10294 genes with data in both H3K79me2 generated in this manuscript and Shao et al. datasets were grouped into quintiles based on the gain of H3K79me2 signal in Slow versus WT from TSS to TES^24^ . Elongation rates were transformed with log10 scale. p < 2.22e-16 by a Wilcoxon signed-rank test^24^ .

We hypothesized that the elevated H3K79me2 observed in 2i/LIF ESCs could also be due to altered RNAPII dynamics, specifically, the reduced elongation rates. To test this hypothesis, we obtained an ESC line that contained a R749H point mutation in RNAPII, known to cause a slower elongation rate, and propagated the cell line in 2i/LIF conditions^34^. While total RNAPII levels were invariant in wildtype and slow RNAPII conditions, there was a slight decrease in Ser2P, Ser5P, and DOT1L levels (Figure 1C). Nonetheless, H3K79me2 increased by 1.5-fold (Figure 1C). To determine whether specific genomic locations were susceptible to the gain of H3K79me2 under slow RNAPII conditions, we performed chromatin immunoprecipitation followed by sequencing (ChIP-seq). Since H3K79me2 is enriched on the gene body, we stratified all genes into quintiles of largest to smallest gain in H3K79me2 enrichment in the slow RNAPII ESC line compared to wild type ESCs (Figure 1D). The genes in the bottom quintile (largest gain) were those with the highest pre-existing levels of H3K79me2 and RNAPII occupancy (Figure 1E). Notably, these genes also tended to have higher transcriptional elongation rates, implying that slow mutation does not deposit H3K79me2 at de novo locations but amplifies the endogenous H3K79me2 signal (Figure 1F). DOT1L interacts with actively transcribing RNAPII and mutant RNAPII has a longer residence time over the gene body and therefore accumulates more H3K79me2 at more actively transcribing genes.

Collectively, our results suggest that both the increased DOT1L abundance, and the slower RNAPII elongation can contribute to the increased H3K79 methylation observed in 2i/LIF ESC. This mechanism may contribute to the heightened H3K79me2 levels observed in 2i/LIF ESCs.

### DOT1L overexpression alters RNAPII occupancy irrespective of catalytic activity

We next wanted to determine if elevating DOT1L or H3K79me2 in serum/LIF ESCs could affect RNAPII enrichment pattern. We generated ESC lines that integrated either wild-type (DOT1L-WT) or a catalytically inactive mutant (DOT1L-CM) version of FLAG-tagged DOT1L into a single location in the genome under the control of a doxycycline-inducible promoter^35^ . Upon acute doxycycline (dox) induction for 24 hours, we observed an approximately 8-fold increase in DOT1L protein expression in three independent clones of both DOT1L-WT and DOT1L-CM cell lines (Figure 2A). As expected, overexpression of DOT1L-WT led to a ∼2.5-fold increase in H3K79me2, while the DOT1L-CM did not alter H3K79me2 levels (Figure 2A).

**Figure 2.**
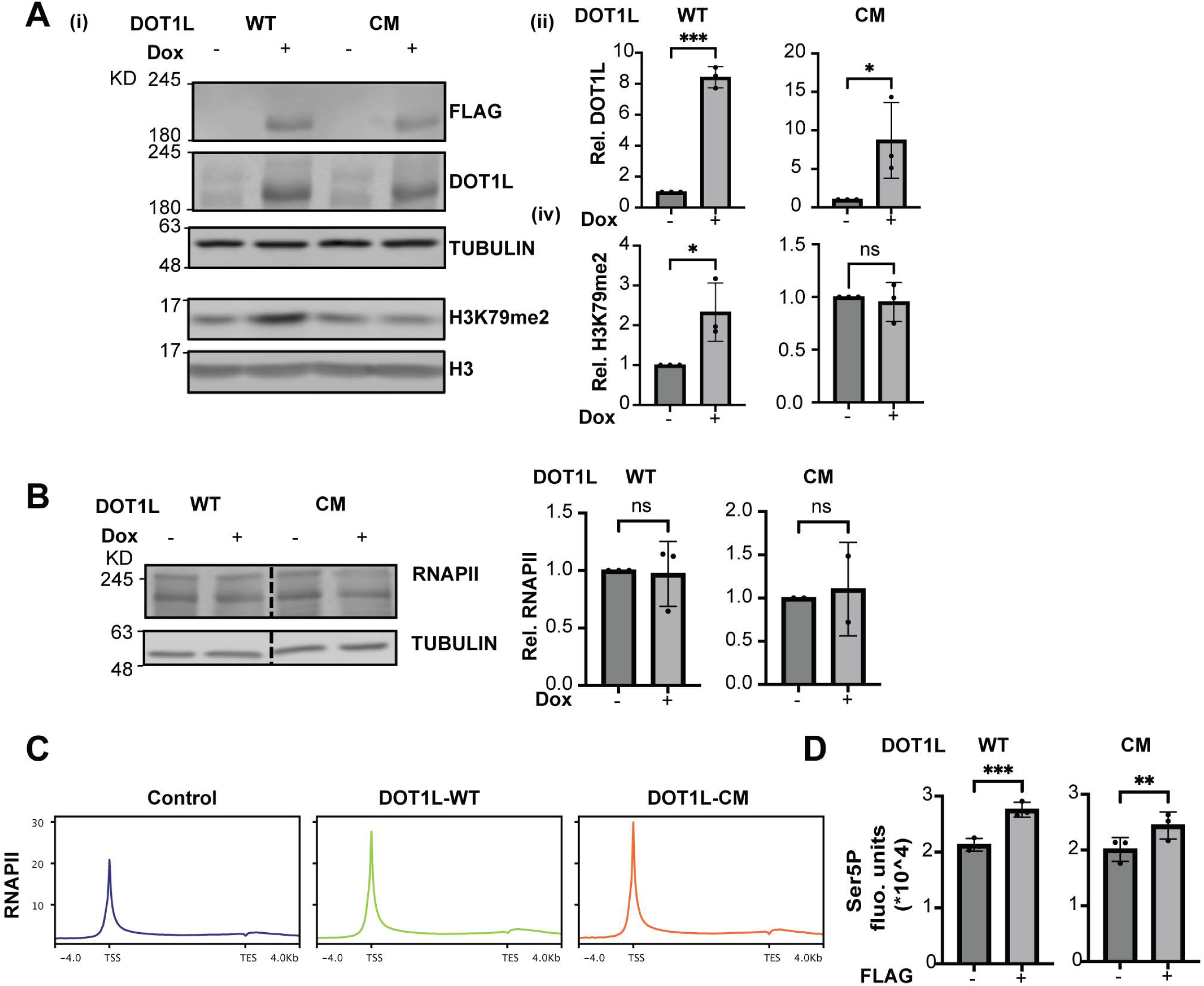
**Inducible overexpression of DOT1L increases RNAPII at TSS independent of catalytic activity.** (A) (i) Western blots of FLAG-tagged DOT1L, total DOT1L, and H3K79me2 after 24 hr dox (Dox) induction in WT or CM ESCs. TUBULIN and H3 serve as loading controls respectively. (ii) Quantification of relative DOT1L levels normalized to TUBULIN uninduced controls set to 1 (mean ± SD, *** = p < 0.001, * = p < 0.05, ratio paired two-tailed t test, n = 3). (iii) Quantification of relative H3K79me2 normalized to total H3 (mean ± SD, * = p < 0.05, ns = not significant, paired two-tailed t test, n = 3). (B) Western blots of RNAPII in Dox-induced (+) WT and CM ESCs compared to uninduced controls (-), with TUBULIN loading control. Bar graphs show relative RNAPII levels normalized to TUBULIN, uninduced controls set to 1 (mean ± SD, ns = not significant, ratio paired two-tailed t test, n = 3). (C) DOT1L overexpression increases RNAPII Ser5P phosphorylation without altering total RNAPII distribution. Metagene profiles of RNAPII ChIP-seq signal around transcription start sites (TSS) and transcription end sites (TES) in control, WT, and CM overexpression conditions. (D) Quantification of relative RNAPII Ser5P fluorescence intensity in FLAG-positive (+) compared to FLAG-negative (-) after FLAG-DOT1L-WT or -CM DOT1L induction. FLAG-negative (-) population set to 1. (mean ± SD, ** = p < 0.01, *** = p < 0.001, paired two-tailed t test, n = 3).

Since slower RNAPII could increase H3K79me2, using these DOT1L ESC lines, we could determine whether there was a reciprocal regulation of RNAPII. Therefore, we performed ChIP-seq of RNAPII after DOT1L induction. DOT1L overexpression resulted in a global increase in RNAPII occupancy of genes, especially at the transcription start sites (TSSs) (Figure 2C). Interestingly, this effect was observed for both DOT1L-WT and DOT1L-CM, suggesting that the influence of DOT1L on RNAPII localization at the TSS is independent of its catalytic activity. We confirmed that there was an increase in H3K79me2 at three genic locations that gained RNAPII only upon induction of DOT1L-WT by ChIP-qPCR (Supplementary Figure 1B). This increase in RNAPII at the TSS was not due to an increase in RNAPII protein levels (Figure 2B). However, reflecting the greater enrichment of RNAPII at the TSS, we observed an increase in the initiation-specific Ser5P form of RNAPII in DOT1L-induced cells (Figure 2D).

RNAPII patterns can vary among transcribed genes with varying enrichments at TSS, the transcription end site (TES), and the pause site. We grouped genes based on RNAPII occupancy patterns at and found that DOT1L overexpression reinforces the pre-existing RNAPII profiles (Supplementary Figure 1A). For example, genes that exhibited high RNAPII levels across gene bodies under control conditions (clusters 1 and 2) showed increased RNAPII occupancy throughout gene bodies upon DOT1L overexpression. In contrast, genes that were characterized by high levels of paused RNAPII at the TSS displayed RNAPII increases primarily at the TSS. Notably, the magnitude of RNAPII accumulation at the TSS was comparable across genes regardless of their patterning (Supplementary Figure 1A). Therefore, a specific pattern was not favored for RNAPII stalling by DOT1L overexpression.

### DOT1L overexpression alters RNAPII occupancy in a context-dependent manner

To identify genomic and chromatin features associated with the extent of RNAPII increase upon DOT1L overexpression, we stratified genes into quintiles of increased RNAPII occupancy at the TSS (Figure 3A). The median increase in RNAPII enrichment across the quintiles ranged from 1.2 to 1.6-fold over serum ESC. For the three middle quintiles, the increase in RNAPII enrichment corresponded with the initial levels of RNAPII (Figure 3B). Befitting the RNAPII enrichment, the middle three quintiles also displayed greater absolute expression as well as higher H3K79me2 enrichment of the corresponding genes (Figure 3C and 3E)^36^ . Interestingly, both the first and fifth quintile had lower initial RNAPII, H3K79me2, slower elongation rates, and were lowly expressed genes (Figure 3B-E)^24^ .

**Figure 3.**
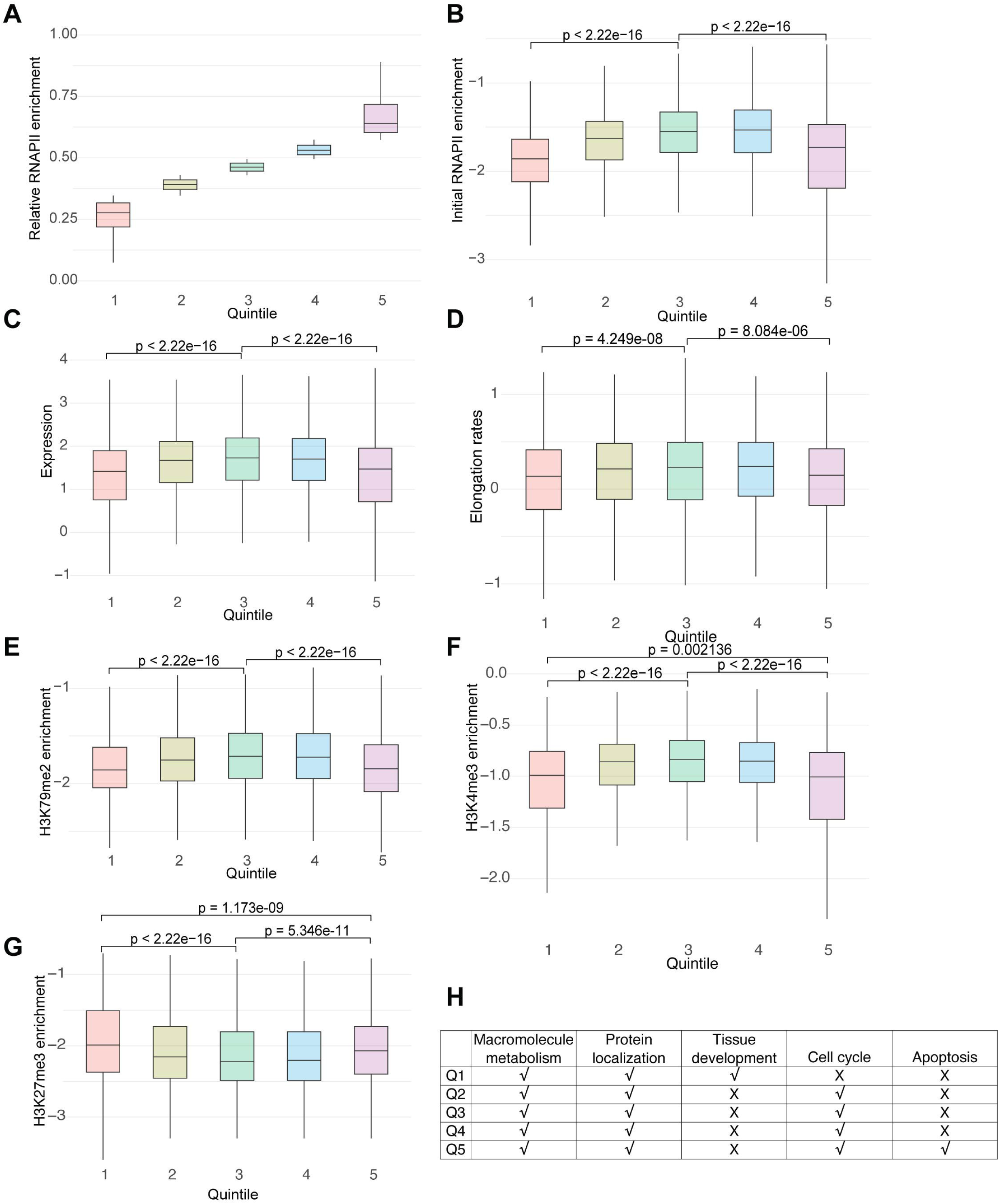
**Gene features associated with the magnitude of RNAPII enrichment.** (A) Box plot of relative RNAPII in DOT1L overexpression versus control, genes are quantified from −30 to +300 bp of TSS and grouped into quintiles. (B-G) Box plot of initial RNAPII in control (B) Absolute expression level in serum/LIF ESCs (Bulut-Karslioglu et al.)^36^ (C) Elongation rates (Shao et al.)^24^ (D) H3K79me2 enrichment from TSS to TES (E) H3K4me3 enrichment (Marks et al.)^16^ from −2kb to +2kb of TSS (F) H3K27me3 enrichment from −2kb to +2kb of TSS at genes within quintiles from (A). All data was normalized to quantified region length and log10 transformed. Statistical significance for box plots was determined by a Wilcoxon signed-rank test. (H) Gene Ontology analysis of Quintiles from (A). Selected enriched GO terms per quintile.

We determined the enrichment of two other histone modifications, H3K4me3 and H3K27me3, using published data^16^ . While H3K4me3 enrichment was higher in quintile 1 than quintile 5, quintile 1 genes had a more significant enrichment of H3K27 methylation within 2 kb of the TSS than quintile 5 (Figure 2F and 2G). While genes in all quintiles functioned in housekeeping functions such as metabolism and intracellular transport, gene ontology analysis demonstrated that quintile 1 and quintile 5 were uniquely enriched for developmental process functions, and cell cycle and apoptosis, respectively (Figure 2H).

Given this increase in RNAPII, we next determined if there was a consequence on gene expression. Bulk RNA-seq analysis revealed that overexpression of DOT1L—either wild-type or catalytic mutant—did not result in significant changes in the expression of any genes, aside from DOT1L itself (Supplementary Figure 2A). To assess whether DOT1L-induced RNAPII accumulation affects nascent transcription, we selected three representative genes that showed increased RNAPII enrichment at their TSSs and measured nascent transcript levels using intronic qPCR (Supplementary Figure 1B and 2B). These analyses revealed no substantial changes in nascent transcription. Thus, the immediate effect of RNAPII accumulation upon DOT1L overexpression does not translate into increased transcriptional initiation or elongation to affect steady-state expression.

Taken together, highly expressed genes in serum/LIF ESCs can gain RNAPII proportional to existing RNAPII, H3K79me2, and absolute expression levels. By contrast, lowly expressed genes that are H3K27me3-enriched and lineage-specific are more resistant to RNAPII gain as compared to those that function in cell cycle/ apoptosis.

### H3K27me3 restricts DOT1L-induced RNAPII accumulation at bivalent genes

H3K27me3 is deposited by the Polycomb complex that contains a methyltransferase subunit EZH2 and a subunit that propagates the methyltransferase, EED. In a prior study, an EED knockout ESC line propagated in 2i conditions was sufficient to decrease H3K79me2 levels, suggesting a crosstalk between these histone modifications^23^ .

To determine whether DOT1L overexpression reciprocally influences the balance of H3K27me3 in serum/LIF ESCs, we examined global H3K27me3 levels in ESC lines pulsed with DOT1L-WT or DOT1L-CM. We found that neither DOT1L-WT nor DOT1L-CM altered global H3K27me3 levels, suggesting that DOT1L does not immediately cause a change in H3K27me3 deposition (Figure 4A).

**Figure 4.**
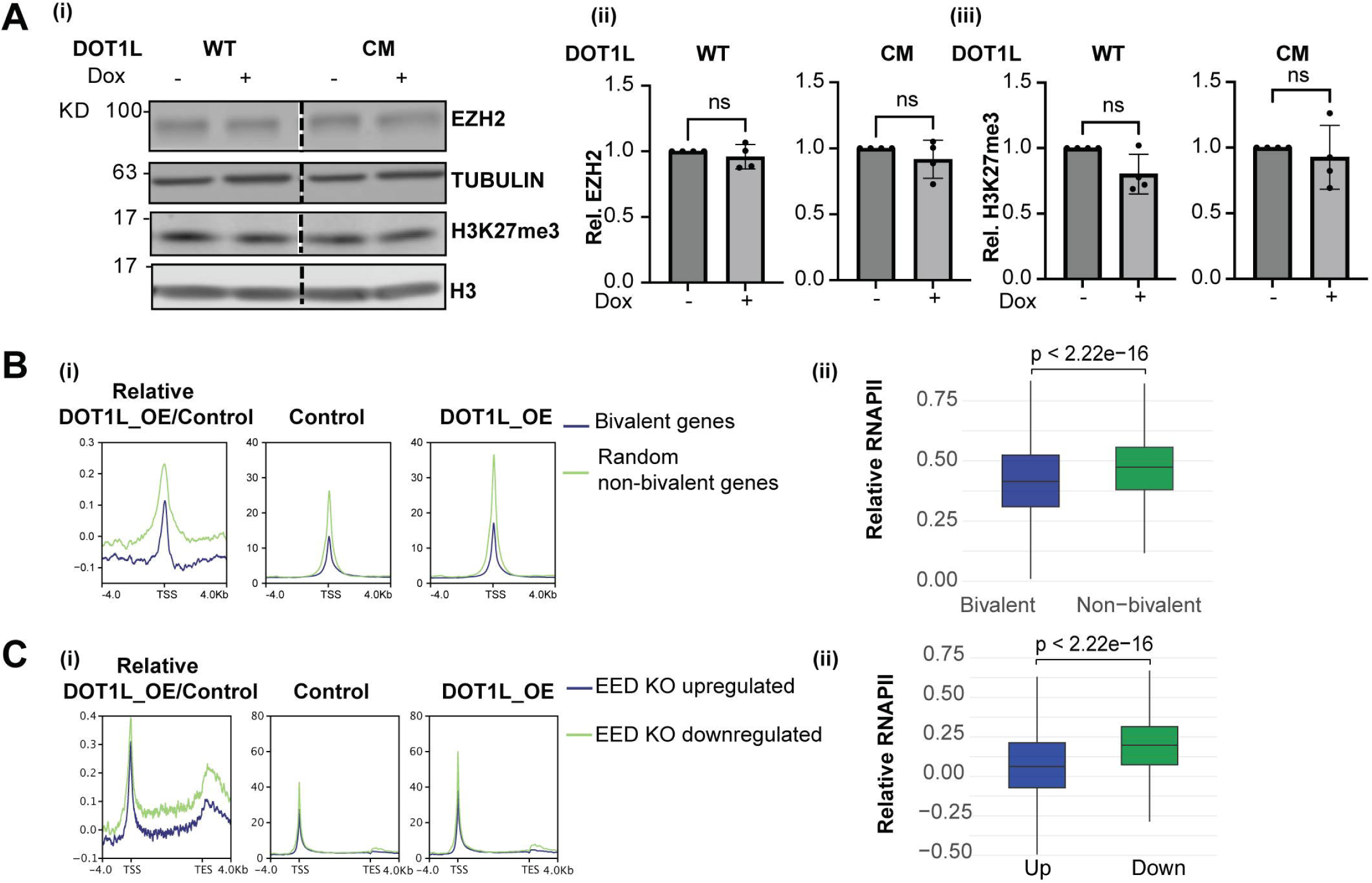
**DOT1L overexpression mediated RNAPII enrichment is mitigated at PRC2-repressed genes.** (A) (i) Western blot of EZH2 with TUBULIN control and H3K27me3 with H3 control in DOT1L-WT or -CM overexpression serum/LIF ESCs with (+) or without (–) Dox induction for 24 h. (ii–iii) Quantification of EZH2 (ii) and H3K27me3 (iii) normalized to TUBULIN or H3 respectively (mean ± SD; ns, not significant by ratio paired two–tailed t-test; n=3). (B) (i) Metaplots showing relative RNAPII enrichment in DOT1L overexpression vs control (left), absolute RNAPII levels in control (middle) and DOT1L-OE ESCs (right) at bivalent (blue) vs random non-bivalent (green) genes. (ii) Boxplot of relative RNAPII log2 fold change in DOT1L-OE vs control for 3641 bivalent vs 9262 non-bivalent genes (p < 2.22e-16, Wilcoxon rank-sum test). (C) (i) Metaplots showing relative RNAPII enrichment in DOT1L overexpression vs control (left), absolute RNAPII in control (middle), and DOT1L-OE ESCs (right) for genes upregulated (green) or downregulated (blue) upon EED knockout in 2i/LIF ESCs (Mierlo et al.)^23^ . (ii) Boxplot of relative RNAPII log2 fold change in DOT1L-OE vs control for EED KO upregulated vs downregulated genes (p < 2.22e-16, Wilcoxon rank-sum test).

Serum/LIF ESCs maintain a transcriptionally poised chromatin landscape by depositing both the activating mark H3K4me3 and the repressive mark H3K27me3 at promoters^16,20,21^ . This bivalent configuration enables repression of lineage-specifying genes while allowing for rapid activation in response to differentiation signals. Bivalency is highly reduced on 2i conditions due to a drop in both H3K4me3 and H3K27me3^16,22,23^ . We then asked if the gain in RNAPII upon DOT1L induction was related to bivalency. We compared RNAPII changes at bivalent genes to those at a randomly sampled background gene set(Figure 4B). Bivalent genes gained less RNAPII supporting the conclusion that the repressive influence of H3K27me3 limits DOT1L-mediated RNAPII recruitment.

To further test the functional relationship between H3K27me3 and DOT1L-mediated RNAPII dynamics, we examined a published dataset of gene expression changes in EED knockout 2i/LIF ESC^23^ . Genes that were upregulated upon EED deletion—i.e., genes that were previously repressed by H3K27me3—were more responsive to DOT1L overexpression than those that were downregulated (Figure 4C). This result suggests that DOT1L preferentially affects RNAPII occupancy at genes not actively repressed by H3K27me3.

Together, these findings demonstrate that H3K27me3 limits the ability of DOT1L to promote RNAPII accumulation, particularly at bivalent genes, and that the chromatin context modulates the transcriptional responsiveness to DOT1L activity.

### DOT1L overexpression recapitulates 2i/LIF state RNAPII dynamics

Previous studies have shown that ESCs cultured in 2i/LIF conditions exhibit increased RNAPII pausing compared to serum-grown ESCs. The increased RNAPII signal at the TSS observed upon DOT1L overexpression mimics 2i-associated RNAPII gain. To evaluate the extent of this resemblance, we compared DOT1L-induced RNAPII distribution changes with published RNAPII ChIP-seq data from 2i and serum LIF ESCs^16^ . We performed k-means clustering of the RNAPII enrichment patterns based on the difference between both 2i vs serum LIF ESCs and DOT1L overexpression in serum/LIF ESCs (Figure 5A).

**Figure 5.**
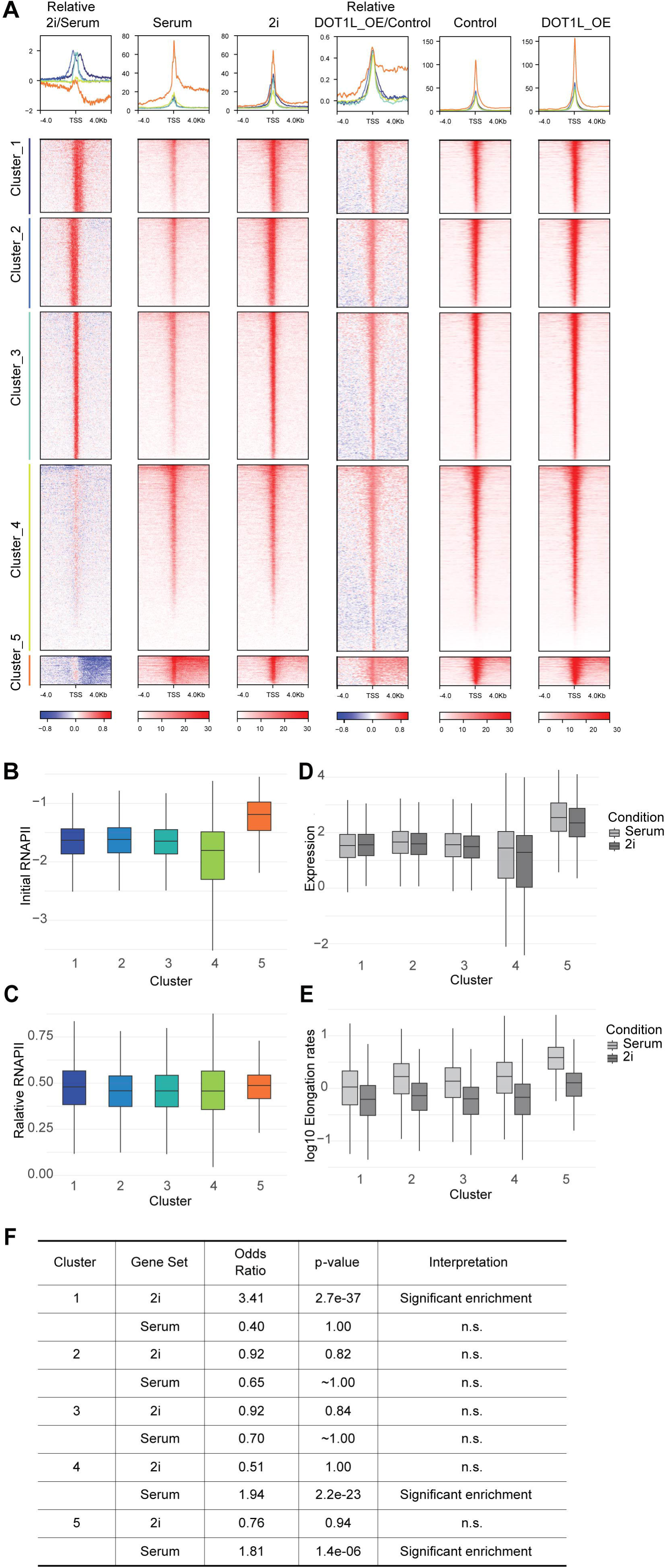
**Clustering of RNAPII occupancy reveals distinct gene groups with differential responses to pluripotent states and DOT1L overexpression.** (A) Metagene profiles and heatmaps of RNAPII occupancy across gene clusters. Top panels show metaplots of RNAPII ChIP-seq signals ±4 kb from TSS and TES in serum versus 2i ESCs (left three columns)^16^ and control versus DOT1L overexpression (right three columns). Relative fold change profiles are shown for serum vs 2i (far left) and DOT1LOE vs control (middle) and divided into five distinct clusters with different response patterns. (B) RNAPII levels (log 10 scale) in control serum/LIF ESCs across clusters in (A). (C) RNAPII signal fold change (log2 scale) in each cluster after DOT1L_OE relative to control conditions across clusters in (A). (D) Gene expression (log10 scale) levels in serum (light grey) versus 2i (dark grey) ESCs from Bulut-Karslioglu et al. RNA-seq data across clusters in (A). (E) RNAPII elongation rates (log10 scale) in serum (light grey) versus 2i (dark grey) ESCs from Shao et al. datasets across clusters in (A). (F) Fisher’s exact test to assess the enrichment of genes upregulated in 2i or serum-passaged ESCs across RNAPII occupancy clusters (from Figure 5A) using published RNA-seq data^16^ .

Clusters 1 through 3 had an equivalent amount and more TSS-biased pattern of RNAPII enrichment in both serum and 2i LIF ESCs (Figure 5A-C). By contrast, Cluster 5 had more RNAPII in the gene body in serum ESC as compared to a TSS-restricted pattern in 2i/LIF ESCs (Figure 5A). Cluster 5 genes were more highly expressed, and the highest RNAPII elongation rates was observed in both serum and 2i conditions (Figure 5D and 5E). Notably, Cluster 5 genes exhibited the most substantial reduction in elongation rate in serum vs 2i conditions (Figure 5E). To further characterize the genes in these clusters, we included a gene expression dataset that performed RNA-seq analysis on serum vs 2i passaged ESCs^36^ . Cluster 1 had a significant enrichment for 2i-enriched genes, while

Cluster 5 had a significant enrichment for serum-enriched genes (Figure 5F). For clusters 1 through 3, the pulse of DOT1L overexpression enabled additional RNAPII occupancy, albeit not to the same levels as 2i/LIF ESCs (Figure 5A). Therefore, the increased DOT1L protein levels observed in 2i ESC may be an important contributor to the RNAPII dynamics for these genes, especially in Cluster 1. For Cluster 5, additional factors present in 2i/LIF medium may be essential for the resolution of the serum-like RNAPII pattern.

### DOT1L overexpression influences NANOG homogeneity but does not induce transcriptional quiescence

Gene regulation in both serum and 2i LIF ESCs is dependent on the core pluripotency factors OCT4/SOX2 and NANOG. However, *Nanog* is heterogeneously expressed in serum/LIF conditions^21^ . When serum/LIF ESCs are transferred to 2i/LIF medium, the NANOG protein distribution in an ESC colony becomes more homogenous within 24 hours accompanied by a minor increase in *Nanog* expression.

Upon DOT1L induction, we observed that cells with higher levels of FLAG expression exhibited more homogeneous NANOG expression (Figure 6A and Supplementary Figure 3A). This occurred in both DOT-1L WT and DOT1L-CM cell lines. In neither condition was there an immediate increase in *Nanog* mRNA expression (Figure 6B). Thus, DOT1L overexpression in serum/LIF recapitulates NANOG homogeneity in 2i/LIF ESCs through non-transcriptional mechanisms.

**Figure 6.**
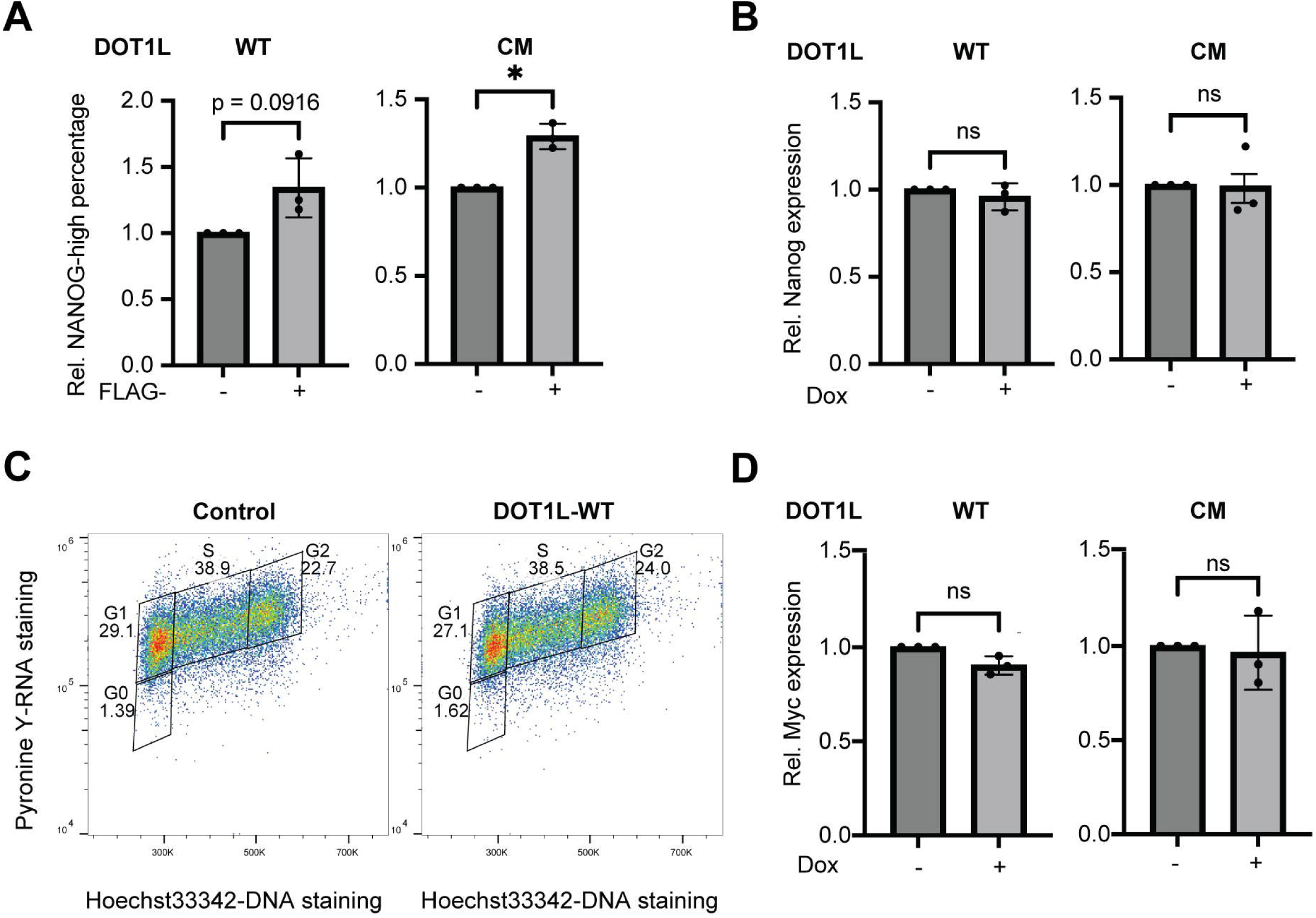
**DOT1L overexpression increases homogeneity of NANOG** (A) Percentage of NANOG-high cells in wild-type (WT) and catalytic mutant (CM) DOT1L-overexpressing serum/LIF ESCs after 24h Dox with FLAG-High (+) or -Low (-) populations. FLAG-Low populations were set to 1. Populations were gated as described in Supplementary Figure 3A. Bars represent median ± SD (n = 3 biological replicates; *p < 0.05; ns: not significant; ratio two-tailed paired t-test). (B) Relative *Nanog* mRNA expression measured by RT-qPCR in DOT1L-WT or -CM ESCs with (+) or without (-) 24h Dox induction (n = 3 biological replicates; ns: not significant; ratio two-tailed paired t-test). Controls were set to 1. (C) Quiescence assay by Hoechst33342 DNA content and Pyronin Y RNA staining showing gating of distinct cell cycle states (G0, G1, S, G2) in Control (left) and DOT1L-WT overexpressing 2i/LIF ESCs with (right) or without (left) 24h Dox induction. Representative flow cytometry plots are shown. (D) Relative c-*Myc* mRNA expression measured by RT-qPCR in WT and CM DOT1L-overexpressing ESCs with (+) or without (-) 24h Dox induction. Bars represent mean ± SD (n = 3 biological replicates; ns: not significant; paired two-tailed t-test). Controls were set to 1.

We next considered whether DOT1L overexpression may influence transcriptional pausing or quiescence, phenomena closely associated with the pluripotent state. The transcription factor MYC is highly expressed in serum/LIF ESCs but significantly reduced in 2i/LIF conditions. MYC is known to promote global transcriptional amplification and RNAPII pause-release. Inhibition of MYC induces a quiescent state marked by cell cycle exit and reduced transcriptional activity, particularly in 2i/LIF ESCs ^37^ .

We hypothesized that if DOT1L overexpression enhances RNAPII pausing, it might promote a quiescent-like transcriptional state, analogous to the effect of MYC inhibition. However, our data showed that DOT1L overexpression alone is not sufficient to increase quiescence in 2i/LIF ESCs (Figure 6C and 6D). Despite its impact on RNAPII localization and NANOG expression homogeneity, DOT1L does not recapitulate the full transcriptional shutdown or cell cycle exit associated with MYC suppression.

## Discussion

In summary, acute DOT1L overexpression induces changes in RNAPII occupancy and reinforces aspects of the ground state of pluripotency—such as uniform NANOG expression—without altering global transcription or promoting transcriptional quiescence in ESCs. We observed that the catalytic mutant of DOT1L, while incapable of increasing H3K79me2 levels, still promoted RNAPII recruitment at TSS, reinforcing the conclusion that DOT1L-mediated RNAPII regulation is partially catalytic-independent. Additionally, NANOG expression heterogeneity decreased in FLAG-DOT1L-overexpressing cells, mimicking the more homogeneous expression profile typical of 2i/LIF ESCs, though this did not translate into MYC-inhibition-like quiescence.

We discovered that 2i/LIF ESCs exhibit distinct RNAPII dynamics, characterized by a prolonged residence time that facilitates the accumulation of DOT1L and a subsequent increase in H3K79 methylation. Reciprocally, our results demonstrate that DOT1L protein can recruit RNAPII to chromatin, forming a positive feedback loop that enhances RNAPII occupancy in ESCs. Notably, this recruitment occurs independently of the catalytic activity of DOT1L, indicating a non-enzymatic role in RNAPII deposition.

Our findings are consistent with previous reports showing loss of RNAPII at TSS upon DOT1L depletion in mouse ESCs^29^ . However, neither DOT1L overexpression nor depletion significantly alters steady-state or nascent transcription levels, within the short time course we examined. This suggests that changes in RNAPII occupancy do not immediately translate into altered transcriptional output.

DOT1L overexpression does not produce uniform changes in RNAPII accumulation. DOT1L-induced RNAPII accumulation is mitigated at genes marked with H3K27me3. These results point to H3K27me3 as a potential repressive mark that blocks DOT1L-mediated RNAPII recruitment, especially at bivalent genes.

The absence of nascent transcriptional changes following DOT1L overexpression likely reflects the early time point (24 hours) chosen to capture direct, rather than adaptive, effects of DOT1L modulation. In future studies, quantitative ChIP-seq using exogenous spike-in chromatin will be essential to more accurately assess global RNAPII redistribution, and genome-wide nascent transcription assays such as TT-seq or GRO-seq will help identify direct transcriptional outputs.

To investigate DOT1L’s immediate effects on RNAPII and chromatin, rapid degradation systems such as PROTACs or degron tags will be valuable tools. In leukemia models, DOT1L inhibition causes H3K27me3 to spread into oncogenes, and in MLL-AF4 leukemias, H3K79me2/3 supports maintenance of H3K27ac at enhancers, underscoring the epigenetic interplay between DOT1L and Polycom-related pathways^35,38^ .

In 2i/LIF ESCs, the global loss of H3K27me3 via EED deletion reduces H3K79me2, highlighting a functional connection between these modifications^23^ . Moreover, co-inhibition of EZH2 and DOT1L has been shown to promote neuronal maturation and block B-cell lymphoma, suggesting that DOT1L and H3K27me3 operate in a coordinated, and sometimes antagonistic, manner in development and cancer^39,40^ .

Finally, based on our results, DOT1L can be added to a growing list of histone modification complexes that are being rediscovered as modulating RNAPII pause release or elongation. The DPY30 subunit of the MLL H3K4 methyltransferase complex, when acutely degraded, causes a decrease in H3K4me3 and an increase in RNAPII pausing^41^ . In fact, we find that the genes that have the largest increase in RNAPII in the gene body upon DPY30 depletion also exhibited increased gene body RNAPII upon DOT1L overexpression (Supplementary Figure 3B and C). This shared response suggests a possible convergence of regulatory mechanisms between DOT1L activity and H3K4me3-dependent control of RNAPII dynamics.

## Materials and Methods

### Cell isolation and culture

R749H mutant E14 ESCs were a kind gift from Cáceres lab^34^ .

Feeder fibroblasts were expanded for three passages and irradiated with 9 krad. ESCs (serum conditions) were cultured on gelatinized dishes with serum/LIF medium (knockout DMEM, 15% FBS, 1× nonessential amino acids, 1× GlutaMAX, 1× penicillin/streptomycin, 4 μl/525 ml of 2-mercaptoethanol, and leukemia inhibitory factor). V6.5 ESCs were grown on irradiated feeder mouse embryonic fibroblasts and passaged with 0.25% trypsin every 1 or 2 days. For 2i/LIF state, ESCs grown in serum/LIF were passaged at least 4 times on 0.1% gelatin-coated plates in modified 2i/LIF medium: 50% DMEM/F12 + N-2 supplement + insulin (12.5 mg/liter) + progesterone (0.01 mg/liter), 50% Neurobasal medium + B-27 minus vitamin A, LIF (1000 U/ml), 2-mercaptoethanol (6.3 μl/liter), penicillin-streptomycin, 3 μM CHIR99021, 1uM PD032590, bovine serum albumin (BSA) (0.0025%), and 1% FBS. Wild-type and R749H mutant E14 cells were cultured without feeders in the 2i/LIF media. 2i-ESCs were passaged with TrypLE every 1 or 2 days. Mycoplasma test was performed monthly for all cell lines.

### Inducible DOT1L ESC lines

5′-FLAG–tagged full-length mouse DOT1L wildtype cDNA with a T2A-linked EGFP or catalytic-dead mutated cDNA with a T2A-linked mScarlet-I sequences were cloned into pBS33 respectively, which was transfected with plasmids encoding FLP recombinase through electroporation. The catalytic dead mutant was generated by mutating amino acids 162-164 from GSG to RCR^42^ . Exogenous DOT1L was integrated into the *Col1a1* locus of V6.5, under the control of a doxycycline-inducible promoter. DOT1L expression was induced with doxycycline (2 μg/ml).

### Reverse transcription qPCR

RNA was isolated from cells with the PureLink™ RNA Mini Kit (Invitrogen, 12183025). One microgram was converted to cDNA with qScript (Quanta, 95047), and 7.5 ng of cDNA (based on the original RNA concentration) was used for qPCR analysis in 12μl reactions with SYBR Green (Bio-Rad, 1725124). Relative expression was normalized to *Gapdh*.

### Immunoblot

SUMO buffer was used to lyse cells containing ^1^/4 part I [5% SDS, 0.15 M tris-HCl (pH 6.8), 30% glycerol], ^3^/4 part II [25 mM tris-HCl (pH 8.3), 50 mM NaCl, 0.5% NP-40, 0.5% deoxycholate, 0.1% SDS], and 1× cOmplete protease inhibitors (Roche, 4693132001). Lysates were sonicated with a microtip for 5s twice at 20% amplitude and quantified in the DC Protein Assay Kit II (Bio-Rad, 5000112) against a BSA standard curve. Twenty micrograms of protein was loaded for H3K79me2 blots, 20 μg for RNAPII blots, 10 μg for H3K9ac, and 20 μg for H3K27me3. Protein was transferred to a nitrocellulose membrane for 16hrs at 100mA and blocked in 5% milk/TBS–0.1% Tween 20 for 30 min. Membranes were probed for 1 hour to overnight in primary antibody, including H3K79me2 (1:1000; Active Motif, 39143), H3K9ac (1:1000; Active Motif, 39917), H3K27me3 (1:1000; Cell Signaling, 9733S and Active Motif, 39155), RNAPII RPB1 NTD D8L4Y (1:1000; Cell Signaling, 14958S), total H3 (1:3000; Cell Signaling, 3638S), and α-TUBULIN (1:3000; Cell Signaling, 3873), Ser2P (1:1000; Millipore, 04-1571), Ser5P (1:1000; Millipore, 04-1572), FLAG (1:3000; Sigma-Aldrich, F3165), DOT1L (1:1000, Abcam, ab239358), EZH2 (1:4000; BD Biosciences, 612667), NANOG (1:1000, Cell Signaling, D2A3#8822). Membranes were washed twice in wash buffer (PBS, 0.1% Tween 20) and incubated with secondary antibody for 1 hour. Membranes were washed three times in wash buffer before being imaged with ECL on ImageQuant LAS 4000 or rinsed once more with TBS buffer before being imaged on a LI-COR Odyssey imaging system. Tiff files were quantified with Image Studio Lite V5.2 using the Add Rectangle function.

### Flow Cytometry

Cells were fix and permeabilized for 0.5-1 hour at room temperature with eBioscience™ Foxp3 / Transcription Factor Staining Buffer Set. 30-50 thousand cells per sample were stained with primary antibody for 1 hour, then secondary antibody for half an hour in 1X Permeabilization Buffer. The following antibodies are used: FLAG (1:5000; Sigma-Aldrich, F3165), Ser5P (1:1000; Millipore, 04-1572), and NANOG (1:500, Cell Signaling, 8822T).

Around 10 thousand cells were collected for each sample. Cell cycle analysis was performed as previously decribed^43^.

### ChIP and library

At least 3 million ESCs were used as starting material. The H3K79me2 ChIPs were performed on a separate biological replicate expansion. The RNAPII ChIPs were performed on 2 separate cell clones for either wildtype or catalytic mutant DOT1L overexpression. Cells were trypsinized and fixed with 1% formaldehyde in suspension for 10 min, rotating. Cross-linking was quenched with 0.14 M glycine for 5 min, cells were centrifuged at 3000*g* for 2 min, and pellets were washed three times with cold 1× PBS and stored in −80°C. Cells were resuspended in 1 ml of lysis buffer [1% SDS, 50 mM tris-HCl (pH 8), 20 mM EDTA, 1× cOmplete (Roche, 4693132001) protease inhibitor] and sonicated on a Covaris S220 Focused-ultrasonicator with the following parameters: 15 cycles of 45 s ON (peak 170, duty factor 5, cycles/burst 200), 45 s OFF (rest) in 6° to 8°C degassed water.

Sonicated chromatin was centrifuged at 21,000*g* at 6°C for 10 min, and the supernatant was collected and quantified with the Qubit DNA HS Assay Kit (Thermo Fisher Scientific, Q32854). Chromatin was aliquoted ensuring that the SDS remained in solution and chromatin was diluted to 1:10 in dilution buffer [16.7 mM tris-HCl (pH 8), 0.01% SDS, 1.1% Triton-X, 1.2 mM EDTA, and 167 mM NaCl with 1× cOmplete (Roche, 4693132001) protease inhibitor] before mixing with antibody. H3K79me2 ChIPs started with 7 μg of mouse chromatin with 0.3ug Drosophila spike-in (sonicated for 12 cycles) generated from Drosophila S2 cell line. 13ug of chromatin with 0.51ug of Drosophila spike-in was used in RNAPII ChIPs. 5 μl of the following antibodies was used: H3K79me2 (Active Motif, 39143), RPB1 NTD D8L4Y (Cell Signaling Technology, 14958S).

ChIPs were incubated for 16hrs at 4°C with rotation, and then Dynabeads were added for 2 hours. Dynabeads were pre-prepared as follows: 25 μl of protein A (Thermo Fisher Scientific, 10002D) and 25 μl of protein G (Thermo Fisher Scientific, 10004D) were combined and washed twice in dilution buffer, and then resuspended in an equal volume of original dilution buffer. Antibody-bead complexes were washed twice for 5 min rotating at 4°C in 0.75 ml of each of the following buffers: low salt [50 mM Hepes (pH 7.9), 0.1% SDS, 1% Triton X-100, 0.1% deoxycholate, 1 mM EDTA (pH 8.0), 140 mM NaCl], high salt [50 mM Hepes (pH 7.9), 0.1% SDS, 1% Triton X-100, 0.1% deoxycholate, 1 mM EDTA (pH 8.0), 500 mM NaCl], LiCl [20 mM tris-HCl (pH 8), 0.5% NP-40, 0.5% deoxycholate, 1 mM EDTA (pH 8.0), 250 mM LiCl], and TE [10 mM tris-HCl (pH 8), 1 mM EDTA (pH 8)] using a magnetic rack. Beads were incubated with 150 μl of TE with 2 μl RNase A (10 μg) added and incubated at 37°C for 30 min. Beads with crosslinked chromatin were reversed overnight with 100 μg of proteinase K in 0.2% SDS at 65°C. DNA was purified with phenol-chloroform extraction with phase lock tubes, followed by ethanol precipitation. Airdried DNA was resuspended overnight in 20 μl ultrapure H2O at 65°C.

For sequencing experiments, the resuspended DNA was concentrated to 5 μl with a SpeedVac and used for library preparation with Ovation Ultralow System V2 (NuGEN, 0344) according to the manufacturer’s instructions. All reactions were performed in half the volume, and adapters were diluted 1:5 in water. Libraries were amplified for 12-14 cycles, and the signal was checked by running 10% on a 1.5% agarose gel before the final bead purification. After bead purification, the remaining library was run on a 1.5% agarose gel, and size selection was performed by cutting from 200 to 400 bp. The library was purified with the Monarch® Spin DNA Gel Extraction Kit (New England Biolabs, T1020). Quantity was checked with the DNA HS Assay Kit (Thermo Fisher Scientific, Q32854) and Agilent TapeStation D5000. H3K79me2 libraries in wildtype and slow-mutated E14s were sequenced 1 × 50 on Illumina HiSeq4000 at the NUSeq Core (Northwestern Feinberg School of Medicine). RNAPII libraries were sequenced 2 x 150 on Illumina NovaSeqX plus at Azenta.

### ChIP-seq data processing

For RNAPII ChIP-seq, raw paired-end reads were trimmed to 50bp with Cutadapt-5.1 and aligned to the mm10 or dm6 genome assembly using Bowtie2. Sam files were converted into BAM files with Samtools-1.2 view. Aligned reads were filtered for proper pairing and mapping quality (MAPQ ≥ 20) using Samtools. Duplicates were marked with samtools addreplacerg then removed with Picard-3.4.0 MarkDuplicates. Only reads mapping to canonical chromosomes were retained. Final BAM files were sorted and indexed with Samtools. Signal tracks (BigWig) were generated using deepTools bamCoverage with RPGC normalization and ignoring chrX chrY chrM. Reads were quantified with BEDtools2.0 multicov. Quantification and quintile grouping of relative RNAPII of DOT1L overexpression versus control was performed using the average of log2 fold change of read count in induced divided by control in the 4 biological replicates (two DOT1L-WT and two DOT1L-CM). For heatmaps and grouping with RNAPII results in Figure 2 and 3, 10675 consensus genes with at least 5 reads in any sample and consistent increase of RNAPII in at least three out of four reps of DOT1L overexpression compared to control and the other rep with no less than 0.81-fold change decrease are included. Four uninduced samples and four induced samples are averaged respectively with deeptools bigwigCompare –operation mean to generate averaged bigwig files, -operation log2 – pseudocount 1 to generate compared bigwig files. Heatmaps and metaplots were made with deeptools plotHeatmap and plotProfile functions following computeMatrix.

In the case of H3K79me ChIP-seqs, raw single-end reads were aligned to the mm10 or dm6 genome assembly using Bowtie2. Sorted bam files were generated as RNAPII ChIP-seqs, duplicates and non-canonical chromosomes were retained. Total dm6-mapped reads were used for scaling factors to normalize bigwig files and reads quantification. Two WT samples and two R749H mutant samples are averaged respectively with deeptools bigwigCompare –operation mean to generate averaged bigwig files, -operation log2 – pseudocount 1 to generate compared bigwig files.

In the case of heatmap plots, the parameters were as follows: scale-regions -b 4000 -a 4000 --regionBodyLength 10000. IGV was used to display example ChIP-seq tracks of normalized bigwig files. Reads over genomic coordinates were quantified with BEDtools2.0 multibamCov. Reads were normalized to the length (bp) of the assessed region. H3K27me3 ChIP-seq bigwig files from Mierlo et al. were downloaded as bigwig files^41^ . RNAPII, H3K4me3, H3K27me3 ChIP-seq from Marks et al. are downloaded and aligned to the mm10 genome in the same pipeline as H3K79me2 ChIP-seq analysis^16^ .

Bivalent genes were extracted from the union of 3 reference datasets: Marks et al., Mikkelsen et al., Galonska et al. and overlapped with our gene list ^16,21,44^. The same number of (3641) non-bivalent genes were sampled for comparison. Gene ontology (GO) was performed with DAVID (https://david.ncifcrf.gov/tools.jsp) Functional Annotation Tool, Gene Ontology GOTERM_BP_3 and_4 and GOTERM_MF_3 and _4, GO clusters with enrichment score higher than 3 were selected.

### RNA-seq library and data processing

Mouse cells were induced for 48 hours in serum/LIF medium with 2ug/mL doxycycline before harvesting. Spike-in Drosophila S2 lysate in TRIZOL was mixed at a ratio of 1:10 of S2 cells to mouse cells before RNA extraction. RNA-seq library was prepared as previously described^45^ . RNA-seq reads were sequenced with Illumina NextSeq 500 for 1 x 75 bp at the NUSeq. Sequenced reads were trimmed using Trimmomatic^46^ with the following parameters: LEADING:3 TRAILING:3 CROP:100 HEADCROP:10 SLIDINGWINDOW:4:15 MINLEN:36. RNA-seq reads were mapped to dm6 and mm9 genomes with STAR_aligner −2.7.8^47^ and raw mapped Drosophila reads were used for scaling factors using MedianNorm function in EBseq and applied to mouse genome analysis for calling differentially expressed genes^46^. Transcripts per million (TPM) for each gene were generated with RSEM-1.2.4^48^ alignment using parameters with -bowtie-m 200 --bowtie-n 2 --forward-prob 0.5 --seed-length 28 for making the scatter plot.^49^ .

### Primers used in this study

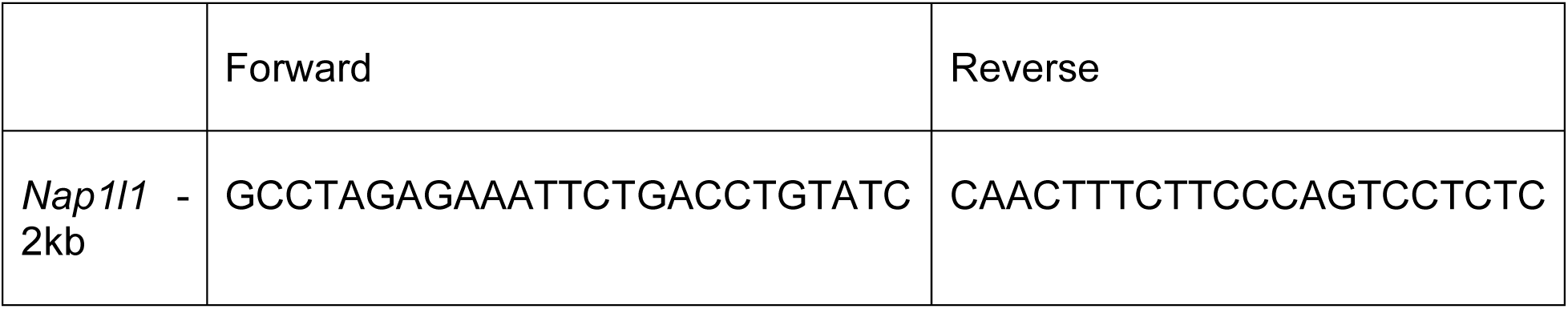
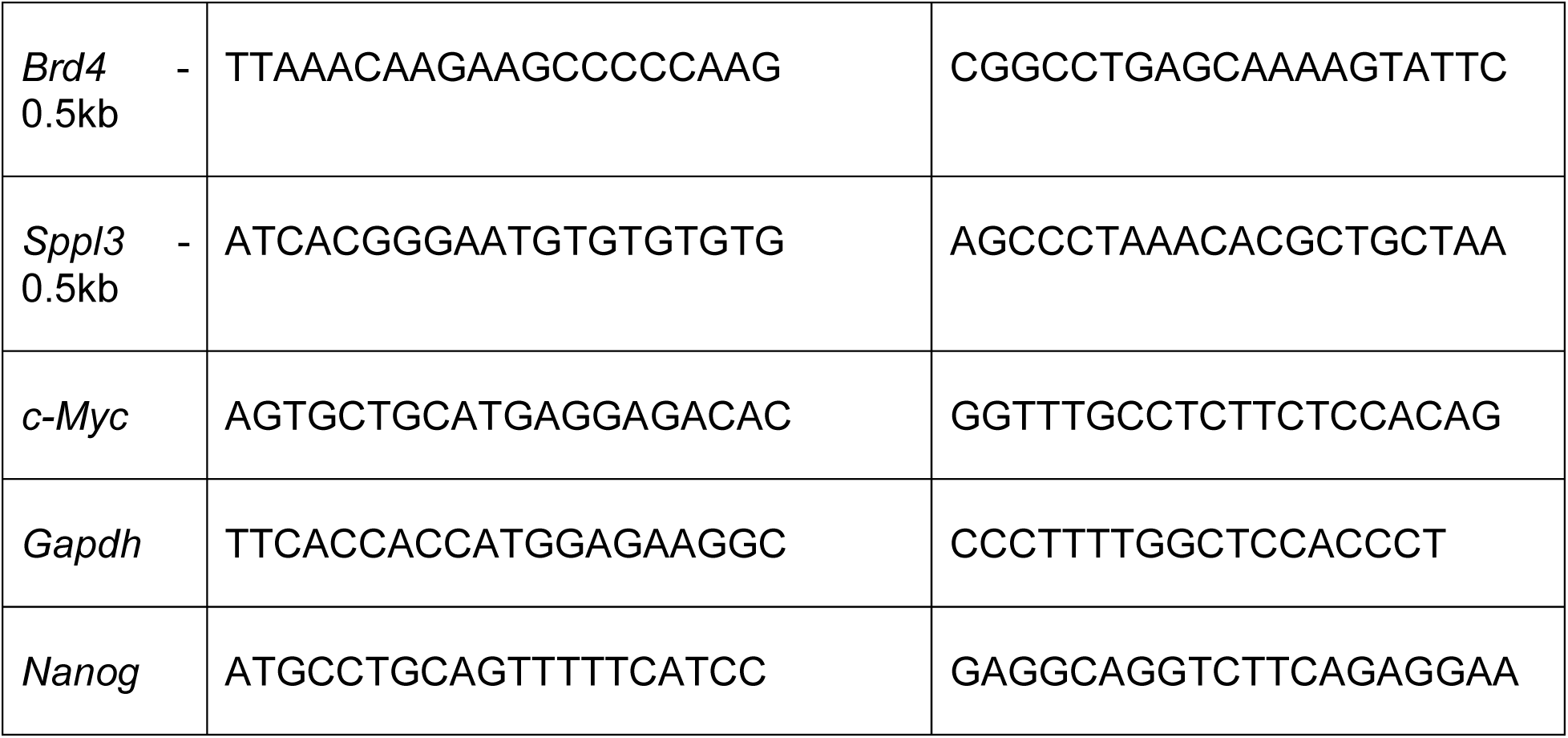

