## Supplementary Figure for "DOT1L induces RNAPII accumulation independent of its catalytic activity in pluripotent stem cells"

### Supplementary Figure 1

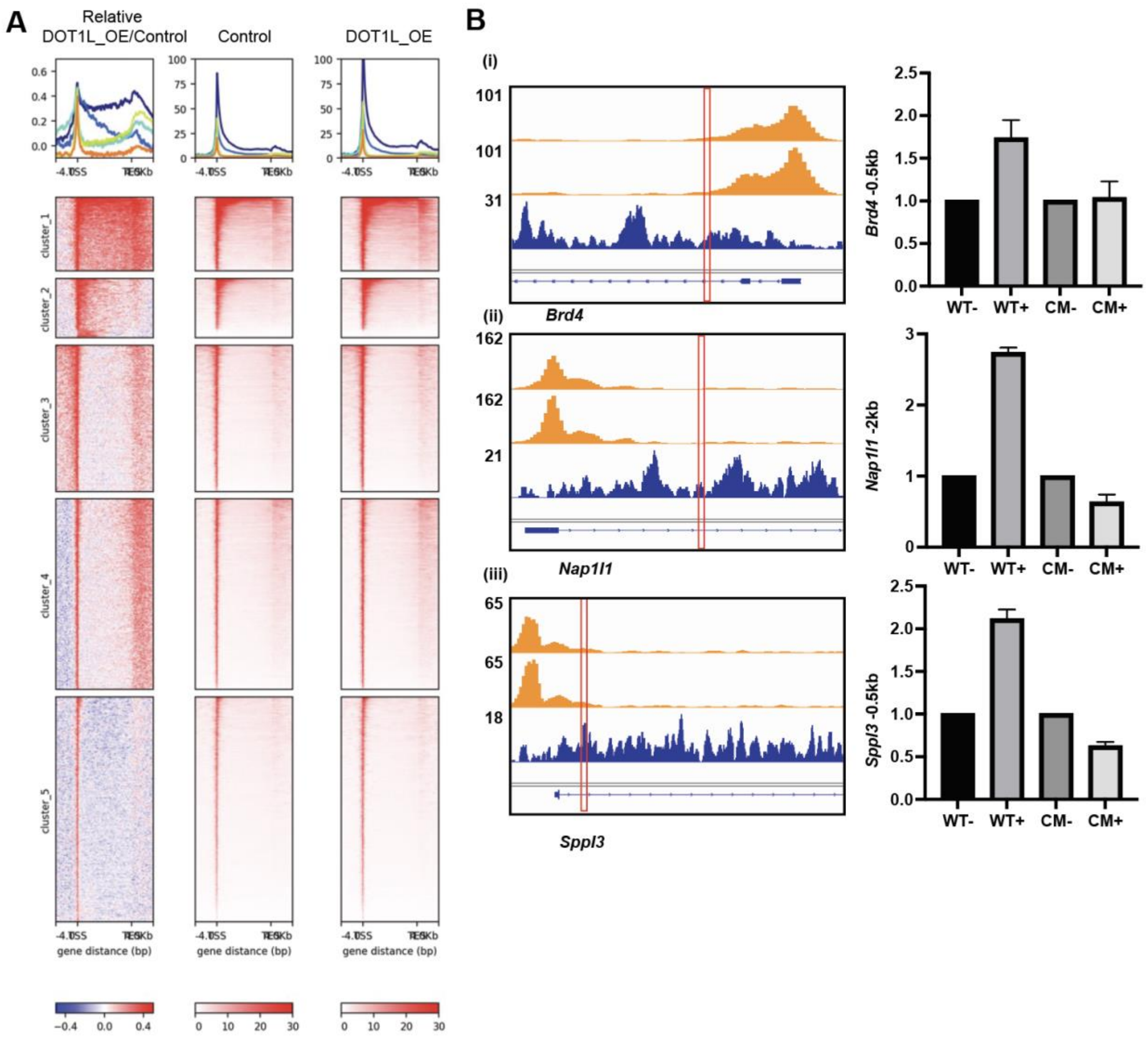


**Supplementary Figure 1. DOT1L overexpression alters RNAPII distribution and affects H3K79me2 at selected loci.**

(A) Metaplots (top) show RNAPII ChIP-seq signals across TSS to TES ± 4 kb for relative fold change (left), control (middle), DOT1L overexpression (right). Corresponding heatmaps (below) were separated into five clusters based on distinct response patterns.

(B) Genome browser tracks and quantification of RNAPII and H3K79me2 occupancy at selected representative genes.

(i–iii) Browser tracks of RNAPII ChIP-seq signal (orange) and H3K79me2 enrichment (blue) at the loci of *Brd4* (i), *Nap1l1* (ii), and *Sppl3* (iii). Red vertical lines indicate the locations of ChIP-PCR primers. Bar graphs on the right show normalized H3K79me2 signal in DOT1L-WT ESCs without (WT−), or with dox induction (WT+), DOT1L-CM without (CM−), and with dox induction (CM+).  n =2.

**Supplementary Figure 2**

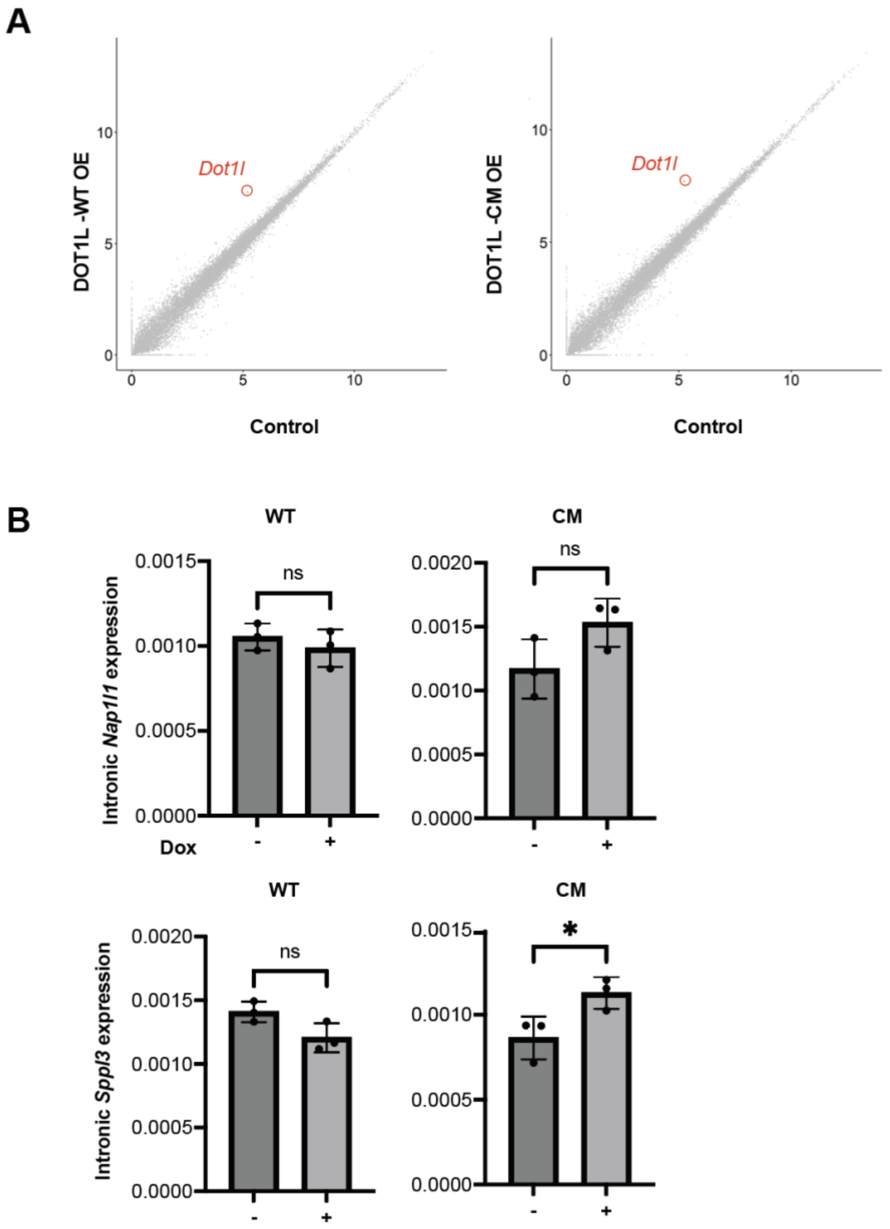


**Supplementary Figure 2. Transcriptomic effects of DOT1L overexpression.**

(A) Scatterplots comparing gene expression (log10-transformed TPM+1) between DOT1L overexpression (WT-OE, left; CM-OE, right) and control ESCs. *Dot1l* is highlighted in red.

(B) Quantification of intronic nascent transcript levels for *Nap1l1* (top) and *Sppl3* (bottom) by RT-qPCR in DOT1L-WT or -CM overexpression ESCs with (+) or without (−) Dox induction for 24 hours. Mean ± SD shown; statistical significance determined by paired two-tailed t-test (ns = not significant, *$p < 0.05$).

**Supplementary Figure 3**

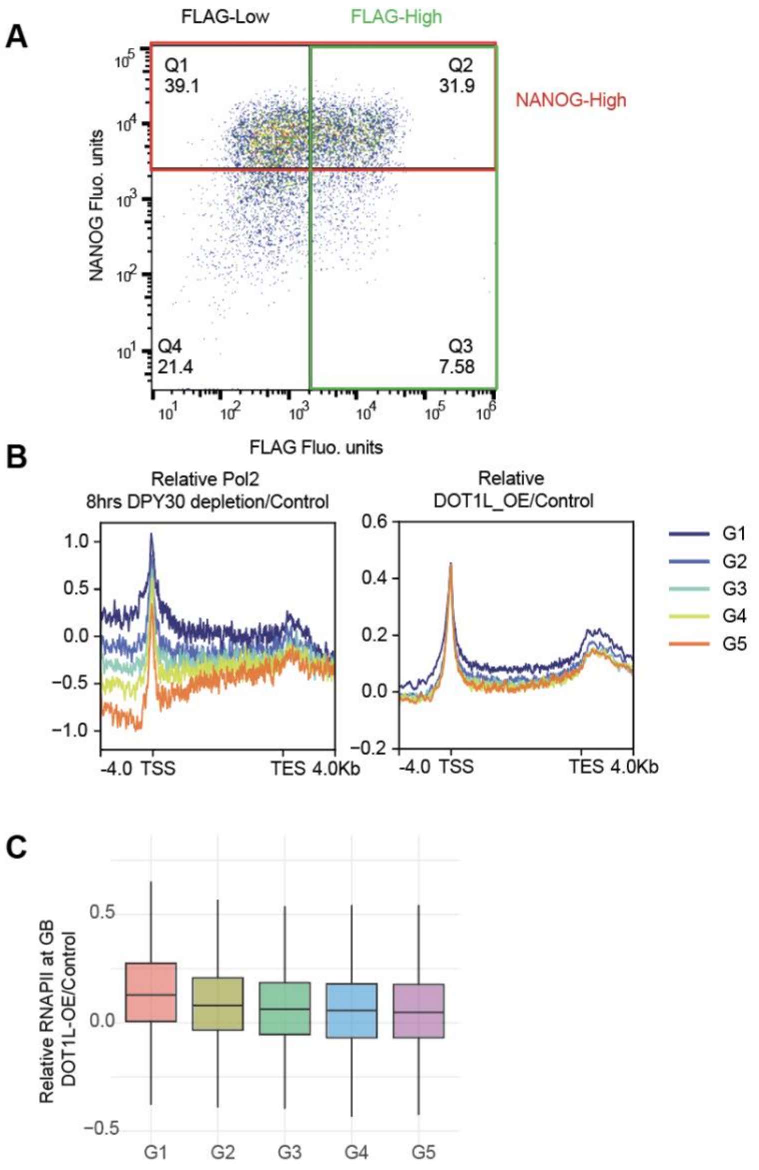


**Supplementary Figure 3. Gating strategy for NANOG-high and FLAG-high cells and gene grouping for RNAPII analyses.**

(A) Flow cytometry gating strategy used to define NANOG-high and FLAG-high populations in DOT1L-overexpressing serum/LIF ESCs. Quadrants represent FLAG-Low/NANOG-Low (Q4), FLAG-Low/NANOG-High (Q1), FLAG-High/NANOG-Low (Q3), and FLAG-High/NANOG-High (Q2). The percentage of cells within each gate is indicated. NANOG-high gating (red line) was used to quantify NANOG-high cell percentages in FLAG-High: Q2/(Q2+Q3) compared to FLAG-Low: Q1/(Q1+Q4) in Figure 6A.

(B) Metaplots of relative RNAPII enrichment sorted and grouped to 5 gene groups by relative Pol II profiles after 8h DPY30 depletion (left, Helin et al.)[1] and relative RNAPII enrichment signal in DOT1L-overexpressing ESCs versus control (right) in color-matched groups of genes.

(C) Boxplots showing relative RNAPII enrichment across the gene body (GB) in DOT1L-overexpressing ESCs compared to control for gene groups in (B).
